# Suppressed macrophage response to quorum-sensing-active *Streptococcus pyogenes* occurs at the level of the nucleus

**DOI:** 10.1101/2025.02.07.637189

**Authors:** Sam F. Feldstein, Kate M. Rahbari, Trevor R. Leonardo, Suzanne A. Alvernaz, Michael J. Federle

## Abstract

*Streptococcus pyogenes*, or Group A Streptococus (GAS), a significant human pathogen, employs quorum sensing (QS) systems to coordinate its behavior and genetic regulation in order to enhance survival. Our previous research established that one such QS system, the Rgg2/3 system, can suppress macrophage NFκB activity and production of pro-inflammatory cytokines. Yet, the scope of suppression and the mechanism by which it occurs remains unknown. In this study, we used transcriptomic and phosphoproteomic approaches to address these unanswered questions. We found QS-ON GAS broadly suppressed most inflammatory transcriptional pathways including those of NFκB, type I and type II interferon responses, and intracellular stress responses. Yet, we found no alternative transcriptional programs were activated after QS-ON GAS infection. Additionally, phosphoproteomics showed no disruption in typical inflammatory pathways such as those related to NFκB and MAPK activation, which was confirmed by western blotting and translocation assays. Instead, the proteomic data highlighted a potential role for epigenetic mechanisms of inflammatory regulation. To determine if epigenetic regulation was involved in QS-mediated immunomodulation, DNA methylation was measured and studies were performed inhibiting various histone and chromatin modifiers. These studies also showed no difference between QS-ON compared with QS-OFF infected macrophages. These findings expand our understanding of QS-mediated suppression and of GAS virulence strategies that appear to employ unusual methods of restricting inflammation. Uncovering this mechanism will offer invaluable insight into GAS, itself, as well as understudied immunological pathways.

**Importance:** *Streptococcus pyogenes* is a ubiquitous pathogen that causes over 600 million infections every year and 500 thousand to 1 million fatalities. While in developed countries it is generally known to cause mild conditions such as pharyngitis, it can also manifest as severe infections such as necrotizing fasciitis, septic arthritis, and lead to post-infectious sequelae including rheumatic heart disease and glomerulonephritis. Elucidating new mechanisms of virulence in this organism, including how it evades and suppresses immune responses can be critical in understanding its pathogenicity, epidemiology, and identification of novel treatment avenues in this era of multi-drug-resistant bacteria. In this study, we characterize the broad spectrum by which GAS modulates the host innate immune response and begin to uncover host pathways that bacteria can use or inhibit for its survival.

## Introduction

*Streptococcus pyogenes*, also referred to as Group A Streptococcus (GAS), thrives as a ubiquitous human pathogen due to its extensive array of virulence factors. To maximize fitness and competitiveness inside the host, GAS employs various mechanisms to survive against the host immune system. One type of system that GAS can utilize is quorum sensing (QS), which is any regulatory system that enables coordinated genetic expression across a population of bacterial cells in response to microbially produced, inter-cellular signals. Other bacterial species such as *Staphylococcus aureus and Pseudomonas aeruginosa* also use their own QS systems to evade the immune system often via the release of toxins (1,2). In GAS, we previously identified and characterized the highly conserved Rgg2/Rgg3 QS system, which confers it with multiple mechanisms of protection against host defenses. These include lysozyme resistance, biofilm formation, and modulation of the host immune system (3–5).

The Rgg2/3 QS system consists of two transcriptional regulators, Rgg2 and Rgg3, which together coordinate activities to control genetic regulation of the system. Rgg2 functions as the transcriptional activator of the system, whereas Rgg3 is the transcriptional repressor (4). In the unstimulated state, Rgg3 predominates over the system and prevents transcription of regulated genes. Under particular conditions, the pheromone SHP (short hydrophobic peptide) can rise in concentration extracellularly. SHP peptides are imported to the cytosol where they bind to both receptors Rgg2 and Rgg3, which derepresses Rgg3 and activates Rgg2, driving transcription of a relatively small gene regulon (4,6). Targets of the system are *shp* itself, thus incorporating a positive-feedback regulatory loop, *stcA,* which we’ve previously demonstrated promotes biofilm formation and confers the bacteria lysozyme resistance, and the 10-gene *spy49_0450-0460* operon that we are in the process of characterizing (3–5,7). Given the distinct regulatory functions of Rgg2 and Rgg3, mutant strains lacking *rgg3* (*Δrgg3*) produce SHP pheromones constitutively and are therefore QS-active (referred to as ‘QS-ON’), while strains lacking *rgg2* (*Δrgg*2) cannot be stimulated and are thus QS-inactive (referred as ‘QS-OFF’). Wildtype strains grown in a nutrient-rich chemically defined medium (CDM) are found to be in a QS-inactive state due to rapid turnover of SHP, unless exogenous SHP is supplied (≥ 1nM) to the culture that results in robust activation.

We previously determined that QS-ON GAS inhibits macrophage NFκB transcriptional activity as well as the production of cytokines including TNFɑ, IL-6, and IFNβ. We further demonstrated that this suppressive phenotype is dependent on each of the first 9 genes in the *spy49_0450-0460* operon (5). Ongoing work is aimed at characterizing the products and effects of this operon.

The ability of GAS to manipulate immune cell signaling has been incompletely characterized. GAS virulence factors have been most well studied for their ability to inhibit phagocytosis (8–10). A growing number of virulence factors have been shown to degrade host proteins including immunoglobulins (11,12), cytokines (13), and complement components (14). One of the best studied proteins is SpeB, a virulence factor regulated by the RopB QS system widely capable of cleaving both host and bacterial proteins (15–17). However, while many Gram-negative pathogens directly interfere with or suppress host immune signaling or activation, far fewer such virulence factors have been reported in Gram positive bacteria (18,19). *Staphylococcus aureus* secretes the protein extracellular fibrinogen-binding protein (Efb), which binds and disrupts TRAF3 in macrophages, leading to suppression of NFκB and AP1 signaling and the reduction of pro-inflammatory cytokine activation (20). An example that shares possible similarities to the suppression we observe in GAS is seen in *Mycobacteria tuberculosis*, which alters its mycolic acid structure, allowing it to bind the host receptor TREM2 to suppress an immune response (21,22). In GAS, capsular hyaluronic acid has been shown to interact with inhibitory host receptors leading to an attenuated immune response, however, our previous studies confirmed that capsule was not necessary for QS-mediated suppression (5,23).

Whereas our previous studies revealed that GAS utilizes QS-signaling to suppress innate immune responses, the mechanism by which this occurs remains elusive. To address this gap, transcriptomic and proteomic approaches were utilized to globally examine differences in macrophage signaling networks after infection with QS-ON or QS-OFF GAS. We found that QS-ON GAS broadly suppresses transcription of inflammatory genes including those regulated by NFκB, interferons, and intracellular stress responses. Phosphoproteomic experiments showed that typical inflammatory signaling factors were not differentially activated, which was supported by western blots and investigating the translocation of transcription factors. Rather, results indicate epigenetic or other intranuclear mechanisms cause the extensive transcriptional suppression seen. Uncovering this novel suppressive mechanism can help our understanding of the disease development and severity of this ever-present pathogen.

## Results

### QS-ON GAS globally suppresses macrophage inflammatory responses

To explore the scope and scale of the effect in which QS-ON Group A Streptococcus (GAS) suppresses inflammatory responses, we performed bulk RNA sequencing (RNAseq) of infected RAW 264.7 macrophages, collecting RNA from cells infected with Δ*rgg2* GAS (QS-OFF), Δ*rgg3* GAS (QS-ON), or a 1:1 mixture of Δ*rgg2* + Δrgg3 GAS (QS-MIX), or from uninfected cells after 2, 4, and 8 hours. A total of 14,072 genes were identified, with 1,476 genes having an absolute difference between QS-ON and QS-OFF of 2-fold or greater and an adjusted p-value of at most 0.1 when averaged across all three timepoints. Principal component analysis confirmed the three infection conditions formed three distinct clusters at the 4- and 8-hour timepoints, with the QS-ON and QS-MIX clusters closer together than the QS-OFF cluster (Fig. 1A). It should be noted that the QS-ON and QS-MIX clusters were significantly separated from the uninfected and all 2-hour samples, indicating distinct transcriptional profiles from uninfected cells. We hypothesize this is likely because the RNAseq data reveals that macrophages appear to still respond to QS-ON GAS, if still significantly less than to QS-OFF GAS.

**Figure 1.**
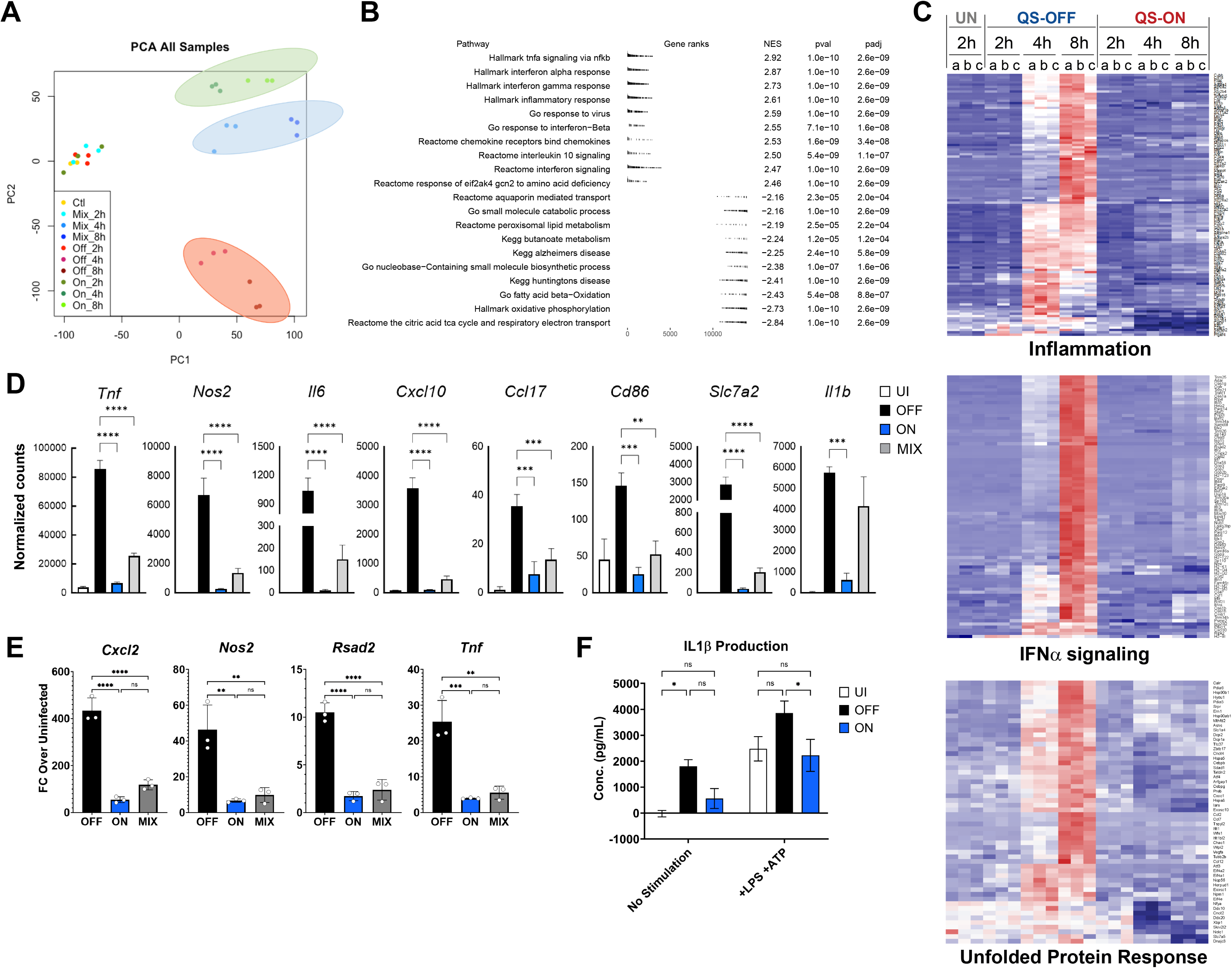
Global suppression of inflammatory genes by QS-ON GAS. A) Principal component analysis showing individual RNA sequencing samples of uninfected and QS- OFF-, QS-ON-, and QS-MIX-infected RAW 264.7 cells at 2-, 4-, and 8-hours post-infection (p.i.) with MOI 40 B) Summary of gene set enrichment analysis comparing QS-OFF- and QS-ON-infected cells. Highlighted are top upregulated and downregulated pathways from GSEA Hallmark, Reactome, GO, and KEGG gene sets. C) Heatmaps comparing Uninfected (UN), QS-OFF-, and QS-ON-infected cells at all timepoints from GSEA Hallmark gene sets of “Inflammation,” “IFNɑ Response,” and “Unfolded Protein Response”. D) Comparison of the normalized read counts from the RNA-seq study from select pro-inflammatory genes. E) Validation of RNA-seq results of select genes by RT-qPCR at 6 hours p.i. with MOI 15 using J774.A macrophages. F) IL1β production from human-derived THP-1 cells infected with QS-OFF or QS-ON GAS. LPS was added 30m p.i., ATP was added 3.5h p.i., and supernatant was collected 4h p.i. with MOI 20. All graphs show mean +SEM. *p<0.05; **p<0.005; ***p<0.001; ****p<0.0001 by one-way ANOVA (D-E) or two-way ANOVA (F) with Tukey’s multiple comparison test.

To identify specific pathways that were differentially regulated after infection with QS-ON compared to QS-OFF GAS, Gene Set Enrichment Analysis (GSEA) was performed. GSEA showed a reduced ability of macrophages to induce a variety of inflammatory pathways, and even some anti-inflammatory pathways, when exposed to QS-ON GAS (Fig. 1B-C). These included pathways related to NFκB signaling, type I and type II interferon signaling, and IL-10 signaling. Inspection of individual genes confirmed that a large number and variety of inflammatory genes were suppressed in QS-ON- and QS-MIX-infected macrophages compared to QS-OFF-Infected (Fig. 1 D). The majority of these inflammatory genes were not significantly upregulated in the QS-OFF condition until 4 hours post-infection. However, closer inspection of individual genes revealed that some were upregulated in QS-OFF-infected cells by 2 hours and remained suppressed in the QS-ON infection, suggesting that suppression occurred early even if the overall effect size was not as strong. These early response genes include *tnf*, *cxcl2*, and *atf3* (Supplemental 1A). Suppression of several genes of interest including *cxcl2*, *tnf*, *pdl1,* and *nos2* were confirmed by RT-qPCR in J774.A cells. (Fig1 E). Additionally, type-I IFN stimulated genes were found to be more suppressed compared to other inflammatory genes, a finding confirmed by RT-qPCR (Supplemental 1B-C). Furthermore, inflammasome suppression by QS-ON GAS was confirmed via ELISA of secreted IL1β in THP-1 cells (Fig1 F). Together, these data indicate a widespread suppression of inflammation that likely impacts a variety of branches of inflammatory programs.

### QS-ON GAS does not induce an alternative transcriptional program in macrophages

We hypothesized that QS-ON GAS might alter macrophage polarization. GSEA analysis indicated that QS-ON-infected macrophages exhibit higher expression of genes in pathways related to fatty acid beta-oxidation and oxidative phosphorylation compared to those infected with QS-OFF GAS (Fig. 1B). However, comparing the fold changes of the genes in these pathways between infected and uninfected cells, the differences are best described as being due to a greater downregulation of these pathways in the QS-OFF-infected condition rather than due to their stimulation by QS-ON bacteria (Supplemental 1D). In fact, of the 2,025 genes more highly expressed in QS-ON-versus QS-OFF-infected cells, only 177 genes were elevated compared to the uninfected control (Fig. 2A). However, these 177 genes are not related by a known common pathway or mechanism by gene ontology (GO) analysis (data not shown). Furthermore, clustering all differentially expressed genes at the 8-hour timepoint across the four experimental conditions revealed 5 distinct clusters (Fig. 2B). Genes in Cluster I correlate to most of the inflammatory genes downregulated after infection with QS-ON or QS-MIX GAS compared to QS-OFF GAS (Supplemental 2A). Clusters II-IV also map well on to known biological functions such as those related to metabolism, chromatin regulation, and cell cycling (Supplemental 2B-D). However, Cluster V, which contains genes that are uniquely upregulated in both the QS-ON and QS-MIX conditions, and would thus most likely inform on a mechanism controlled by QS-ON GAS, did not map to any GO terms. Additionally, as the overall median expression values of genes in cluster V are substantially lower than in the other groups, we suspect these differences may not be biologically relevant (Fig 2C). To determine if the expression levels of any of the genes upregulated in the QS-ON condition compared to QS-OFF appeared biologically relevant, we plotted all the measured genes on an MA plot, which compares the fold change and mean expression value between the two conditions (Fig. 2D). This revealed that genes upregulated in the QS-OFF condition span a wide range of expression, while those upregulated in the QS-ON condition are clustered toward very low expression levels, with many having a normalized read count less than 10. Thus, whereas changes in transcriptional profiles are striking between QS-ON and QS-OFF infections, they did not reveal a clear mechanism for suppression or activation of an alternative transcriptional program.

**Figure 2.**
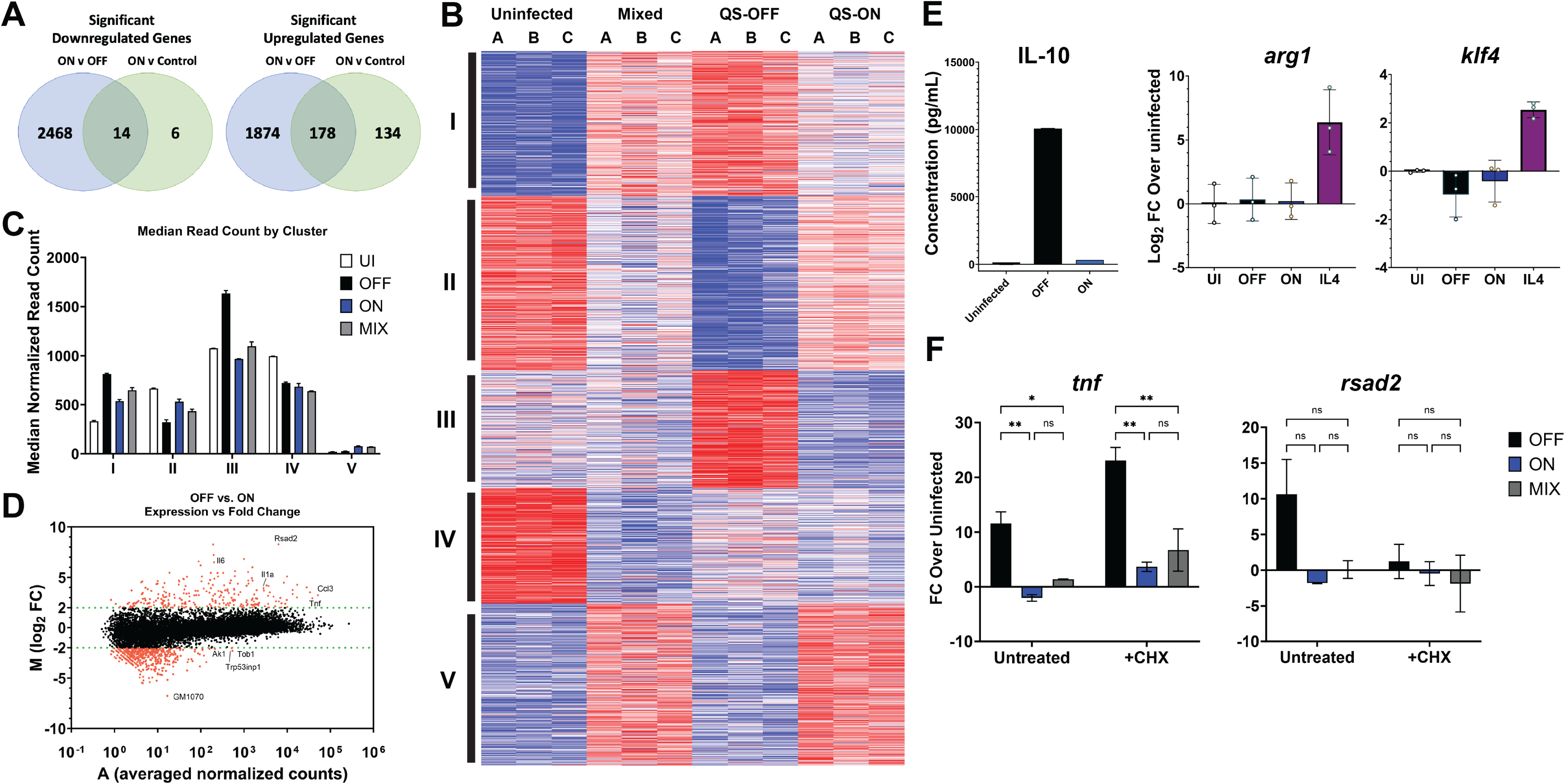
DiGerentially expressed gene clusters do not inform the mechanism of suppression. A) Venn diagrams summarizing the number of genes that are commonly upregulated and downregulated between QS-ON-infected and QS-OFF-infected cells compared to those between QS-ON-infected and uninfected cells when averaged across all timepoints of the RNA-seq analysis. B) Clustering analysis of genes from RNA-seq that showed FDR-adjusted significance between any two conditions at 8h. C) Median normalized read count from RNA-seq analysis for each gene by cluster at 8h. D) MA plot showing the log_2_ fold change between QS-OFF- and QS-ON-infected cells and the average read count between both conditions averaged across all time points. E) IL-10 production in bone marrow derived macrophages cells at 8h post-infection (p.i.) measured by ELISA; gene expression of M2 macrophage markers *arg1 and klf4* in RAW 264.7 cells at 6h p.i measured by RT-qPCR with MOI 15. F) Gene expression of *tnf* (primary response gene) and *rsad2* (secondary response gene) in J774 macrophages at 6-hours p.i. with MOI 20 in the presence or absence of the translation inhibitor cycloheximide (CHX) measured by RT-qPCR. 10 µM CHX was added 15 min prior to infection and remained until cell collection. Graphs show mean +SEM. *p<0.05; **p<0.005 by two-way ANOVA (E) one-way ANOVA (F) with Tukey’s multiple comparison test

Because inducing the secretion of M2-associated cytokines by microbes has been described as a common mechanism to suppress inflammatory responses (24,25), we tested levels of secreted IL-10, an immunosuppressive cytokine, between infection conditions. Consistent with the RNAseq data (Supplemental 1D), IL-10 secretion was highest in the QS-OFF-infected cells, negating the likelihood that M2 polarization accounts for the observed QS-ON-dependent suppression (Fig. 2E). Likewise, common genes associated with alternatively activated macrophages, such as *arg1* and *klf4*, were not upregulated in QS-ON-infected macrophages when measured by RT-qPCR (Fig. 2E). Finally, when macrophages were treated with the translation inhibitor cycloheximide (CHX), QS-ON- and QS-MIX-infected macrophages still exhibited reduced expression of inflammatory genes, indicating the suppression is not caused by any newly transcribed protein induced by QS-ON GAS interaction with the cell (Fig. 2F).

### Phosphoproteome of infected cells suggests an epigenetic mechanism of suppression by QS-ON GAS

Because transcriptomics revealed global immunosuppresion rather than alteration of a specific pathway, we hypothesized that QS-ON GAS might alter inflammatory signaling pathways, which are generally transduced via protein phosphorylation. To test this, we performed a post-translational modification (PTM) scan looking at all detectable phosphorylated proteins 30-minutes post infection of QS-OFF-, QS-ON-, and QS-MIX-infected RAW 264.7 macrophages, compared to steady-state levels of uninfected cells. 20,572 phosphorylation events were detected from 3,393 different proteins. Principal component analysis revealed distinct clusters for the QS-OFF, QS-ON, and QS-MIX conditions (Fig. 3A). Interestingly, the QS-ON infected condition clustered furthest from uninfected condition. Whereas the RNAseq data showed that QS-ON and QS-MIX conditions were more similar to each other, the PTM data showed that QS-OFF and QS-MIX conditions were more similar.

**Figure 3:**
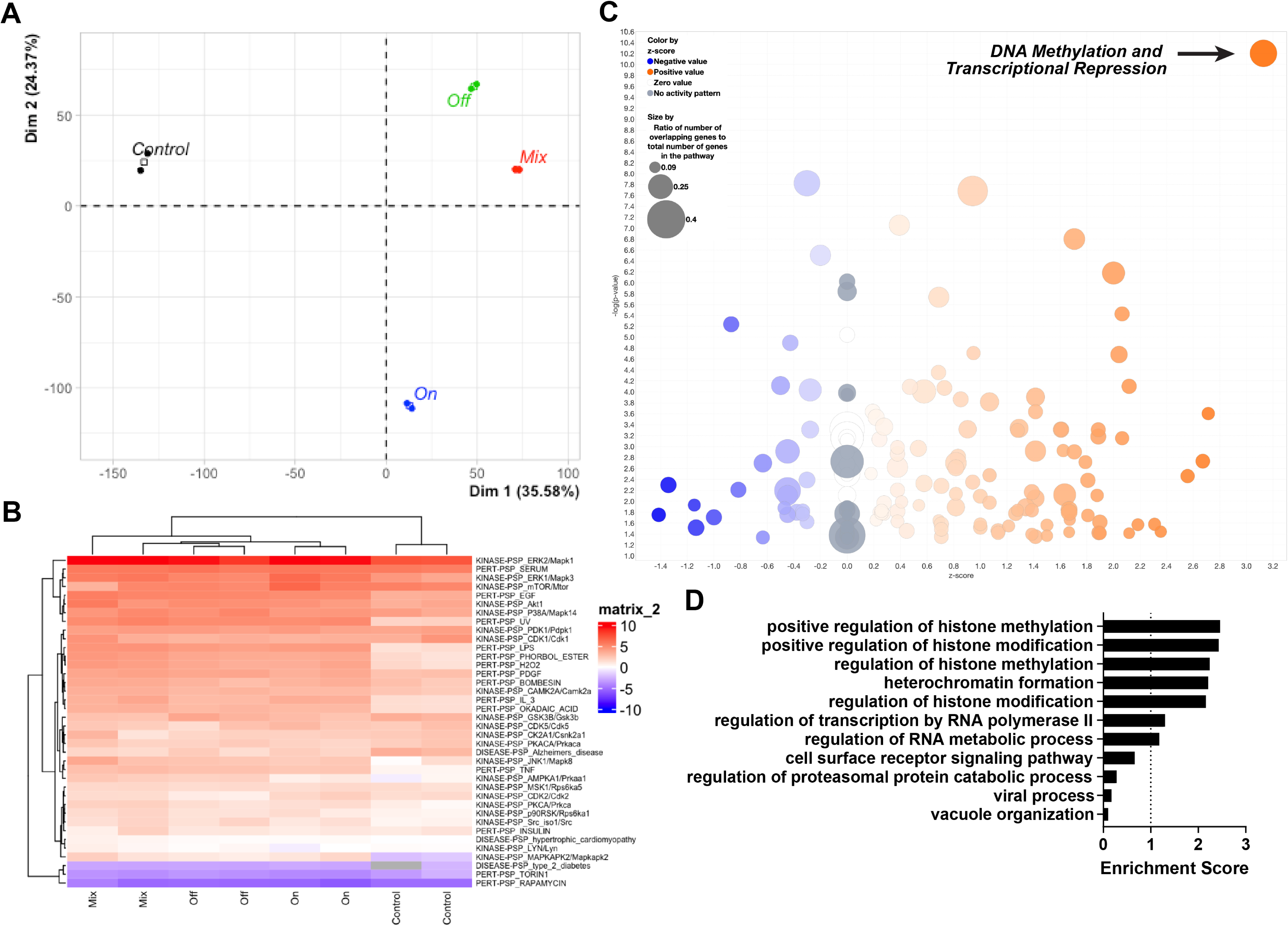
PTM Scan reveal possible epigenetic mechanism of suppression. A) Principal component analysis of post translational modification (PTM) scan measuring the abundances of phosphorylated proteins in uninfected or QS-ON-, QS-OFF-, and QS-MIX-infected RAW 264.7 cells 30m post-infection (p.i.) with MOI 40. B) Heatmap of PTM-signature enrichment analysis (PTM-SEA) showing relative enrichment scores of each sample of each annotated pathway. C) Gene Ontology analysis of PTM scan showing highest enriched pathways upregulated in QS-ON-infected cells compared to QS-OFF-infected D) Ingenuity Pathway Analysis (IPA) of PTM scan depicting its predicted upregulated (orange) and downregulated (blue) pathways in QS-ON-infected cells compared to QS-OFF-infected.

To find pathways that were differentially activated or suppressed between infection conditions via phosphorylation, PTM-Signature Enrichment Analysis (PTM-SEA) was performed using both mouse and human signature databases (human data not shown) (26). Enrichment scores for the PTM-SEA pathways revealed no significant differences between the QS-ON, QS-OFF, and QS-MIX conditions in classical inflammatory or other pathways (Fig. 3B; Supplemental 3A-B). Classical inflammatory pathways and proteins such as those related to TNFɑ, NFκB, and MAPK signaling displayed low levels of phosphorylation in the uninfected condition as expected, while all infected conditions showed similar levels of phosphorylation of proteins in these pathways. Other pathways including those related to CDK2 and GSK3β showed increased and decreased activation, respectively, between QS-ON-infected and QS-OFF-infected cells and between QS-MIX-infected and QS-OFF-infected cells (Supplemental 3B). When performing GO enrichment analysis, comparing phosphorylation levels of proteins in QS-ON- vs. QS-OFF- infected macrophages, several pathways related to epigenetic regulation such as “histone H3-K4 methylation” and “protein deacetylaton”, were enriched (Fig. 3C). This was supported by Ingenuity Pathway Analysis (IPA), in which “DNA Methylation and Transcriptional Repression” was the highest and most significantly upregulated predicted pathway in the QS-ON vs. QS-OFF conditions (Fig. 3D).

Because inflammatory signaling proteins were not differentially phosphorylated, but epigenetic pathways appeared enriched in QS-ON-infected macrophages, we hypothesized that epigenetic or transcriptional regulation within the nucleus may be the point in cellular activation that is impacted by QS-ON-infected cells. To confirm that upstream NFκB activation was not altered between infection conditions, IKK phosphorylation and IkB degradation were evaluated. QS-ON- and QS-MIX-infected macrophages showed similar IKK phosphorylation and higher or equal IkB degradation than QS-OFF-infected macrophages (Fig. 4A). Similar results were observed with or without co-stimulation with LPS (Fig. 4A). Previously, we have shown that NFκB-dependent transcriptional responses induced by LPS and other TLR agonists are suppressed by QS-ON GAS (5). Immunofluorescence microscopy also confirmed that translocation of p65, a typical pro-inflammatory subunit of NFκB, does not exhibit differences in rates of translocation to the nucleus between QS-ON vs QS-OFF conditions (Fig. 4B-C). In fact, none of the five NFκB subunits displayed differences in activation and translocation as determined by an ELISA-based assay (Active Motif) that uses the consensus binding sequence to capture the substrate (Fig. D). These data support the hypothesis that suppression is not occurring at upstream signaling points, but rather within the nucleus after translocation.

**Figure 4:**
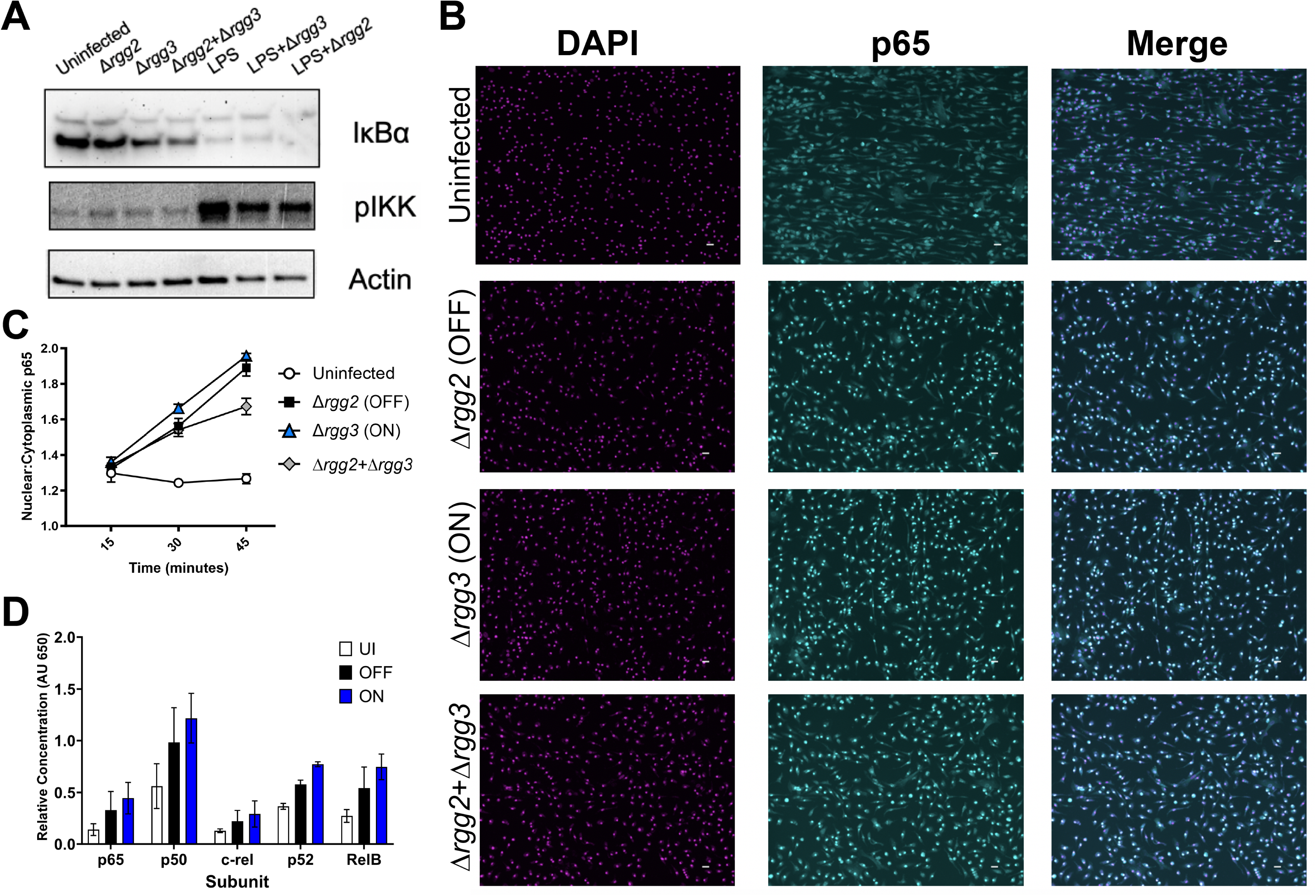
Translocation of NFκB is unaltered between QS-OFF- and QS-ON-infected macrophages. A) Representative western blots of NFκB pathway components IκBɑ and phospho-IKK at 30 min. post-infection (p.i.) with MOI of 40. B) Representative images of immunofluorescence microscopy staining in BMDMs for DAPI (magenta), and p65 (cyan) 45 minutes p.i. with MOI 50 C) Quantification of p65 translocation to the nucleus measured by microscopy. Shown is the mean ratio of mean fluorescence intensity of nuclei compared to cytoplasm as quantified using CellProfiler. Ratios from five fields were averaged. Graph shows mean +SD D) Relative nuclear abundance of each NFκB subunit in differentiated THP-1 cells at 35 min. p.i. with MOI of 20 measured with TransAM NFκB Family Kit from Active Motif; graph shows mean +SEM.

### Direct examination of epigenetic targets and mechanisms does not reveal them as the mechanism of suppression

Given that Ingenuity Pathway Analysis of the PTM data predicted “DNA Methylation and Transcriptional Repression” as the strongest predicted upregulated pathway in QS-ON vs QS-OFF-infected cells, we first examined levels of global DNA methylation of infected cells by an ELISA that measures levels of 5-methylcytosine (5-mC). Given the global suppression observed in the RNAseq data, we hypothesized that if DNA methylation were the mechanism of suppression, it would be measurably evident in such an assay. However, no differences in levels of DNA methylation were observed across any of the infection conditions nor the uninfected control (Fig. 5A).

**Figure 5:**
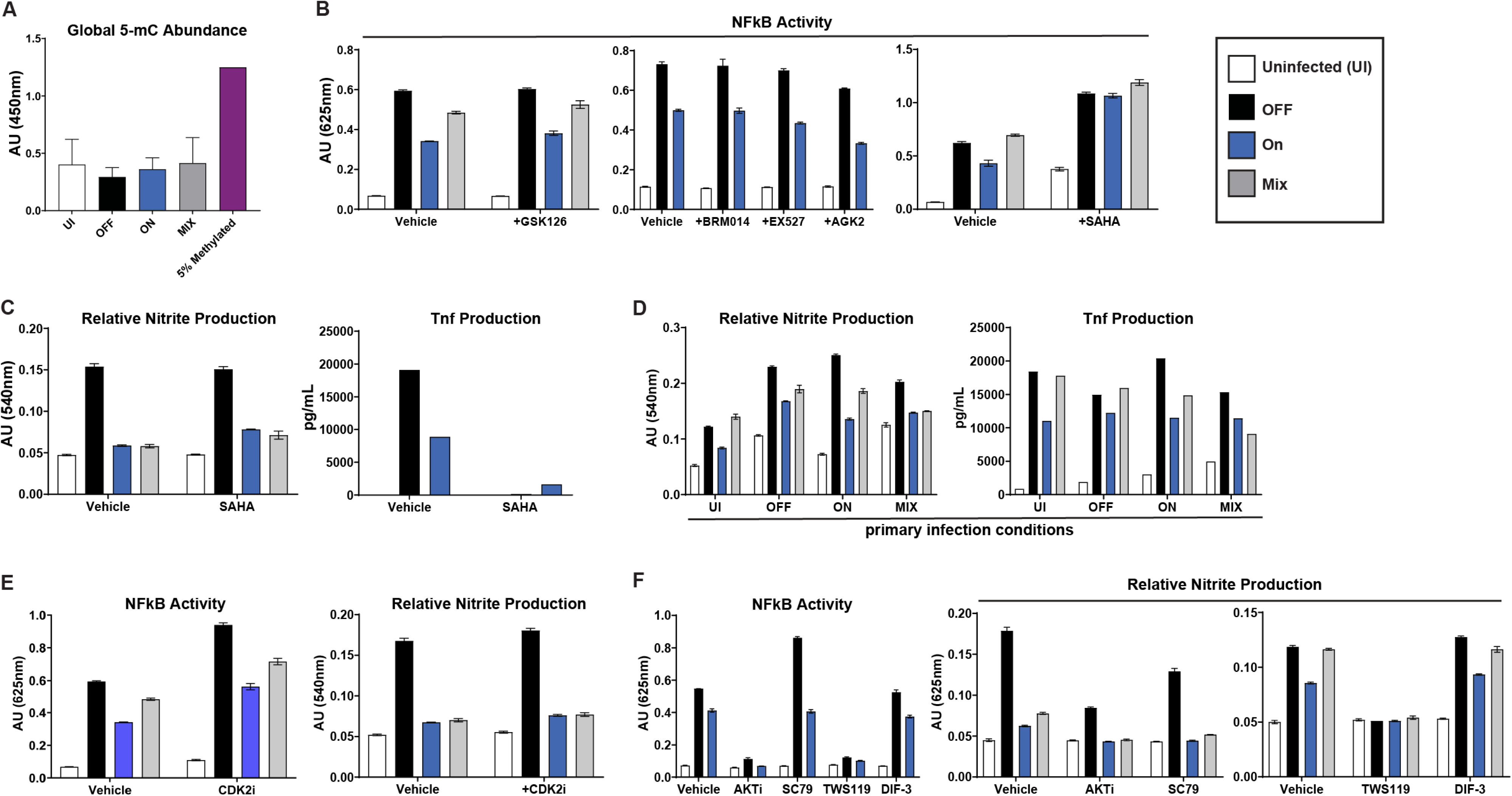
Pharmacological inhibition of epigenetic regulators does not restore inflammatory capacity of macrophages. A) Quantification of global 5-methylcytosine abundance in RAW 264.7 cells 3h post-infection (p.i.) with indicated GAS strains, measured with Epigentek MethylFlash kit. n=2. B) NFκB reporter activities in RAW-Blue cells 18h p.i. with indicated GAS strains treated with DMSO vehicle or 5 µM GSK126, 1 µM BRM014, 10 µM EX527, 5 µM AGK2, or 500 nM SAHA. C) Griess assay and TNFɑ ELISA in RAW 264.7 cells 18h p.i. with indicated GAS strains treated with DMSO vehicle or 500nm SAHA. D) Griess assay and TNFɑ ELISA in RAW 264.7 cells of 18h p.i of a secondary infection. x-axis shows primary infection type with colored bars indicating the secondary infection type. E) NFκB reporter assay and Griess assay 18h p.i. with DMSO vehicle or 2 µM of CDK2 inhibitor II. F) NFκB reporter assay and Griess assay 18h p.i. with DMSO vehicle or 20 µM of AKT1/2 inhibitor, 10 µM of SC79, 7.5 µM of TWS119, or 7.5 µM of DIF-3. All graphs show mean +SEM. All MOI are 10. B- graphs show single experiments representative of 2-4 biological replicates.

Next, to test whether alternative regulation of histones was a mechanism of suppression, several selective inhibitors of histone modifiers were utilized to observe if they blocked the suppression of QS-ON GAS. GSK126 was used to inhibit EZH2, the key catalytic protein of the polycomb repressive complex 2 (PRC2) and shown to be involved in inflammatory regulation (27,28). Suberoylanilide hydroxamic acid (SAHA) is a class I, II, and IV histone deacetylase (HDAC) inhibitor, and is generally shown to have anti-inflammatory properties (29,30). EX-527 and AGK2 were used as sirtuin 1 and sirtuin 2 inhibitors, respectively. Sirtuins are class III HDACs, which have been shown to be targets of bacterial host immune regulation strategies (31,32). Lastly, BRM014 was used to inhibit BRG1, a key component of the SWI/SNF complex, which is involved in chromatin remodeling and shown to be involved in inflammatory processes (33). Neither inhibition of EZH2, nor Sirtuins 1 or 2, nor BRG1 showed the ability rescue full inflammatory capacity of QS-ON-infected macrophages whether measured by TNFɑ production or NFκB reporter activity in RAW Blue cells (Fig. 5B). Although treating cells with SAHA did recover NFκB reporter activity in QS-ON-infected cells (Fig. 5B), this recovery of inflammatory activity was not observed when measuring nitrite and TNFɑ production (Fig 5C). The reduction of TNFɑ in the presence of SAHA is consistent with previous literature regarding its effect on inflammation (29,30). We hypothesize that the reason we observe this effect only in the Raw Blue cells is that the intrachromosomal reporter may be heavily regulated by histone acetylation.

We hypothesized that if QS-ON GAS were inducing an epigenetic change that is causing immune suppression, then that change would be lasting and would be passed on to daughter cells, and thus suppression would remain upon reinfection. This has been demonstrated to occur with other microbes that affect epigenetic regulators. We would expect that after infection with QS-ON, a lasting epigenetic effect would remain so that suppression would still occur upon subsequent infection with QS-OFF GAS. However, this is not what was observed, as after primary infection with QS-ON GAS, a QS-OFF secondary infection could still fully activate the macrophages as measured by Griess assay as well as TNFɑ production (Fig. 5D). Thus, despite the PTM scan analysis, it remains unclear whether epigenetic regulation is truly the mechanism by which QS-ON GAS can suppress macrophages.

We lastly wanted to investigate whether the CDK2 or GSK3β pathways were potentially involved in suppression, given their presence in the PTM-SEA analysis (Fig. S3B). They have been shown to be involved in both inflammatory and epigenetic regulation (34–38). CDK2 Inhibitor II (CDK2i) was used to investigate CDK2, while DIF-3 and TWS119 were used to induce and inhibit GSK3β, respectively. Additionally, because GSK3β is closely tied with the AKT pathway(34), SC79 was used to induce AKT, while Akt1/2 kinase inhibitor (AKTi) was used to inhibit AKT. The CDK2 inhibitor appeared to have no effect on suppression or inflammation (Fig 5E). Both the AKT and GSK3β inhibitors were able to suppress inflammation in all infection conditions, indicating that those are feasible targets for quorum-sensing-dependent suppression (Fig 5E-F). However, if true, we’d expect the inducers to reduce the amount of suppression by QS-ON GAS, which was not observed (Fig 5E-F). Therefore, again, while these were candidates put forth by the PTM analysis, it does not appear quorum-sensing-dependent suppression is occurring by regulation of CDK2, GSK3β, or AKT.

## Discussion

These results show that quorum-sensing-ON Group A Streptococcus (QS-ON GAS) has a global inhibitory effect on inflammatory responses in macrophages, reducing the potency of most, if not all, inflammatory programs. Not only is inflammation significantly attenuated, but no other pathways were seen to be triggered by QS-ON GAS that would account for macrophage inactivation. This is consistent with our findings that the suppression does not seem to be occurring at the point of upstream signaling, but rather it is occurring after translocation of transcription factors within the nucleus at the level of transcription. A post-translational modification (PTM) scan analyzing the phosphoproteome suggests that the suppression is occurring via epigenetic regulation of inflammatory transcription.

Although our previous study characterized the phenomenon of QS-ON as having the ability to actively suppress the TLR-NFκB axis of inflammatory responses, hallmarked by a reduction of key inflammatory markers (i.e IL-6, TNFɑ, IFN), the level of signaling at which activation was blocked, and the scope of suppression was unknown. Additionally, the mechanism of how the suppression occurred was unknown. Here, we expand upon that work by showing the full scale of suppression and demonstrating that suppression occurs after translocation of transcription factors to the nucleus. Although we anticipated QS-ON GAS would prevent activation of NFκB, we saw no differences in phosphorylation of key signaling proteins in NFκB or MAPK/AP1 pathways, and all NFκB subunits translocated to the nucleus at similar rates. Instead, PTM analysis indicates that epigenetic regulation is a possible mode of suppression, which explains why signal transduction pathways promoting inflammation were not altered by QS-ON GAS interaction.

RNAseq datasets did reveal some unanticipated responses to GAS infections. One unexpected pattern related to classical type I interferon (IFN) stimulated genes (ISGs). While classical pro-inflammatory genes such as *tnf*, *nos2*, *il6*, and others exhibit significantly higher expression in QS-OFF vs. QS-ON conditions, they also show significantly higher expression in QS-ON vs. uninfected conditions. Thus, the effect of quorum sensing could be best described as an attenuation of inflammation rather than complete inhibition. However, this is not what is observed with regards to type I ISGs, where complete or near complete inhibition is observed for these genes in QS-ON-infected cells. Classical type I ISGs such as *rsad2*, *mx2*, and the *ifit* genes all have minimal to no upregulation in the QS-ON condition compared to uninfected. We hypothesize the reason for this is that the production of type I IFNs is significantly reduced in QS-ON-infected cells, which we have previously shown by ELISA, and are therefore not producing a sufficient critical concentration to induce production of these ISGs. This may indicate the ultimate benefit received by the bacteria when they activate quorum sensing. While an incomplete reduction in primary response proteins may aid in the survival and persistence of the bacteria, perhaps it is the elimination of the type I IFN response that provides the necessary environment they are seeking. Furthermore, if type I IFN secretion does not reach sufficient concentrations to trigger paracrine signaling *in vitro*, it seems likely that that other attenuated cytokines that communicate to different cells via paracrine or even endocrine signaling would also not be in sufficient concentrations to enact their effects. The effects of type I IFN of GAS fitness and assays measuring cell-to-cell signaling, such as recruitment assays in the context of QS- ON GAS, would be ideal future experiments to perform.

While we have shown that activation of typical inflammatory pathways such those of NFκB and AP1 do not appear to be altered in a QS-ON infection, we are limited by the scope of our post-translational medication scan, which only measured phosphorylated proteins. It is likely that we only see minimal differences in pathways between QS-ON- and QS-OFF-infected macrophages because the signaling is being transduced via alternative modifications. Several inflammatory signaling pathways are induced via signals and switches other than phosphorylation including GTP, cAMP, SUMOylation, ubiquitination, hydroxylation, and others (39–43). However, many of these signals also include a phosphorylation step (such as the cAMP-PKA axis), which were not observed in our PTM scan.

Additionally, while our data strongly suggests an intranuclear mode of inflammatory suppression, we have yet to uncover the particular mechanism. Still, we have confidently ruled out the mechanism being inhibition of upstream signaling molecules in the NFκB and likely AP1 pathways. With these pathways unaltered even up to the point of translocation into the nucleus, the site of inhibition must be within the nucleus. While the epigenetic regulators we investigated did appear to be involved in QS-mediated suppression, we still have not ruled out completely the involvement of another epigenetic protein. This includes those such as ARID4B, which shows significantly higher phosphorylation in the QS-ON- and QS- MIX-infected macrophages. Forthcoming work aims to look at these proteins and regulators as well as direct assessment of histone modifications and chromatin accessibility. Additionally, we have not ruled out the potential role of other intranuclear regulators such as co-activators of transcription, other components necessary for the transcription machinery, or the role of noncoding RNAs. We have equally not eliminated the possibility of post-transcriptional mRNA degradation, a mode of regulation used by pathogens and the host alike (44,45). Lastly, while we have shown that QS-ON GAS is not directly promoting metabolic pathways transcriptionally, metabolic pathways remaining active or silent may also contribute to the observed suppressive phenotype as others have shown that certain metabolic processes may be required to maintain or drive inflammatory states (46–48). Future work will further investigate mentioned alternative modes of signal transduction, possible alternative mechanisms of transcriptional regulation within the nucleus, and finally immunometabolism.

## Methods

### Bacterial Strains

*Streptococcus pyogenes* (Group A Strep, GAS) NZ131 strain was used for all experiments. Δ*rgg2 and Δrgg3* strains were constructed as previously described (4). Starter cultures were used to minimize differences in lag phase between strains and experiments and were created as previously described (5).

### Cell lines

RAW 264.7 murine monocyte/macrophages were cultured in RPMI 1640 (Corning) supplemented with 10% fetal bovine serum (FBS) (BenchMark) and penicillin/streptomycin (P/S) (Corning). THP-1 human monocytes were cultured in identical complete RPMI 1640 media. To differentiate the monocytes into M0 macrophages, cells were treated with 5ng/mL of phorbol 12-myristate 13-acetate (PMA) in complete media for 48h, after which, the cells were washed with PBS and incubated with antibiotic-free and PMA-free media for 24 hours.

J774A.1 murine macrophages were cultured in DMEM (Gibco) supplemented with 10% FBS and P/S. RAW-Blue cells (Invivogen), RAW cells that contain an intrachromosomal insertion of the secreted embryonic alkaline phosphatase gene driven by both NFκB- and AP1-reactive promoters, were cultured in identical complete DMEM with the addition of 200µg/mL of Zeocin every other passage to maintain selection pressure.

### Generation of Bone marrow-derived macrophages

Primary murine bone marrow derived macrophages (BMDMs) were differentiated from bone marrow (BM) cells collected from the femurs and tibiae of six to eight-week-old male or female C57BL/6 (Charles River Laboratories). BM cells were frozen and differentiated at a later date as previously described (5).

### *in vitro* infections

Designated cell lines were seeded into tissue-culture-treated plates (Falcon) in antibiotic-free media and allowed to settle overnight. The following day, GAS strains were grown from frozen starter cultures in CDM to an OD_600_ of 0.5-0.6. The separate strains were normalized by their OD, centrifuged, resuspended in the cell culture medium, then added to the macrophages. The cells were centrifuged at 300 x g for 5 minutes to synchronize bacterial contact with the cells and then incubated for 30 minutes at 37°C with 5% CO_2_ for 30 minutes, unless otherwise noted. After incubation, the wells were washed and replaced with new media containing 100µg/mL of gentamycin (Gibco) to kill any remaining extracellular bacteria.

### RNA-sequencing

RAW 264.7 cells were prepared in RPMI 1640 and infected as described above using a multiplicity of infection (MOI) of 40 CFUs per cell. Cells were infected with Δ*rgg2*, Δ*rgg3*, or MOI of 20 each of Δ*rgg2* + Δ*rgg3*, or bacteria-free media. Infected cells were collected at 2-, 4-, and 8-hours post infection, while the uninfected cells were only collected at 2-hours post infection. At the appropriate timepoint, cells were collected and stored in RNAlater (Invitrogen) at −80°C. Cells were later thawed and their RNA was extracted using RNeasy Mini Kit (Qiagen) and RNase-Free DNase Set (Qiagen) and sent to Novogene for library preparation and sequencing.

At Novogene, RNA was quality control tested via Nanodrop for quantitation, gel electrophoresis for degradation and contamination, and Agilent 2100 Bioanalyzer for integrity and quantitation. mRNA was enriched using oligo(dT) beads and rRNA was removed using Ribo-Zero kit (Illumina). mRNA was randomly fragmented using fragmentation buffer and cDNA synthesized by second strand synthesis using random primers and a custom second-strand synthesis buffer (Illumina). The cDNA was end-repaired, followed by 3’ polyadenylation and sequencing adaptor ligation. The library was then size selected and enriched by PCR. Quality control was performed on the cDNA library via Qubit 2.0 (Thermo) for concentration, Agilent 2100 Bioanalyzer for insert size, and qPCR precise concentration. Qualified libraries were pooled and sequenced paired end at 2 x 150 bp (Illumina).

### RNAseq analysis

Data quality control was performed on FASTQ files using FASTQC and the raw sequencing reads were aligned to the mouse genome (GRCm39) using STAR. Raw sequencing reads of genomic features were enumerated using featureCounts and DESeq2 was used to normalize the counts, and provide differential expression analysis, and pre-ranked test-statistic data. fGSEA algorithm with mouse MSIGDB was used to perform a functional gene set enrichment analysis.

Separately, the featureCounts of uninfected and 8h timepoint samples were normalized and analyzed using edgeR. Genes were then filtered by whether they have an fdr-adjusted p-value <0.05 for at least one pairwise sample comparison. The resulting genes were clustered by k-means clustering, with clustering performed 10 times for a k varying from 2-20. k=5 was chosen by highest reproducibility.

### RT-qPCR

For RNAseq validation and M2 gene investigation, RAW 264.7 cells were prepared in RPMI 1640 and infected as described above using an MOI of 10. Cells were infected with Δ*rgg2*, Δ*rgg3*, or Δ*rgg2* + Δ*rgg3*, or bacteria-free media or stimulated with 100ng/mL of LPS (Sigma) or 20ng/mL of recombinant murine IL-4 (Sigma).

For translation inhibition assays, J774A.1 cells were used. 10 µg/mL of cycloheximide (Sigma) was added 20m before the infection and remained until the cells were harvested.

Cells were collected at 0-, 4-, 6-, and/or 8-hours post infection, and RNA was collected using RNeasy Mini Kit (Qiagen). cDNA conversion was performed using High-Capacity cDNA Reverse Transcription Kit (Applied Biosystems) and qPCR was done using PowerUp SYBR Green (Applied Biosystems). Primers used can be found in Supplemental Table 1.

### Post translational modification scan

RAW 264.7 cells were prepared in RPMI 1640 and infected at an MOI of 40 as described above with Δ*rgg2*, Δ*rgg3*, or Δ*rgg2* + Δ*rgg3*, or bacteria-free media. Cells were collected after 30 minutes and lysed in a 9M urea phospho-protective lysis buffer (9 M sequanol grade urea, 20 mM HEPES, pH 8.0, 1 mM β-glycerophosphate, 1 mM sodium vanadate, 2.5 mM sodium pyrophosphate). Protein concentration of each sample was measured using a DC Protein Assay (Bio-Rad) and Qubit protein assay (Invitrogen), and their concentrations were normalized accordingly. Samples were then sent to Cell Signaling Technology for their PhosphoScan services and analysis. Proteins were digested with trypsin then loaded to a 50 cm x 100 µm PicoFrit column packed with C18 reversed-phase resin. The column was developed with a 150-minute linear gradient in 0.125% formic acid delivered at 280 nL/min. The eluted peptides were lyophilized then phospho-enriched by immobilized metal affinity chromatography using Fe-NTA magnetic beads (Cell Signaling Technology). Phosphorylated peptides were eluted oj the beads with a basic buffer then analyzed by tandem mass spectrometry with parameters optimized by the company. Spectra were evaluated using Comet (49) and the Core platform at Harvard University. Searches were performed against the most recent update (2021) of the Uniprot *Mus musculus* database with mass accuracy of +/- 5ppm for precursor ions and 0.02 Daltons for product ions. Results were filtered with mass accuracy of +/- 5 ppm on precursor ions and presence of the intended motif then further filtered to a 1% false discovery rate.

### PTM Analysis

Any measured protein that did not meet the following quality control checks was removed before analysis: maximum Abundance greater than 1,000,000, maximum coefficient of variance less than 0.5, number of distinct peptides that map to the protein greater than 1. PTM-SEA was then performed using genepattern.com, using their default settings comparing the dataset to both the mouse and human PTM signature databases. Enrichment scores for each annotated pathway of each sample were averaged by experimental group and their differences in means were compared using ANOVA followed by a Tukey post-hoc test.

The dataset was also analyzed using Ingenuity Pathway Analysis (IPA) and by biological process gene ontology (GO) analysis using pantherdb.com. Here, only proteins that exhibited at least a 2.5-fold-change between any two groups were analyzed. Default analysis settings were used for both analyses.

### Western Blot

RAW 264.7 cells were prepared in RPMI 1640 and infected as described above with Δ*rgg2*, Δ*rgg3*, Δ*rgg2* + Δ*rgg3* GAS, or bacteria-free media. Selected wells were also stimulated with 100ng/mL of LPS (Sigma). Cells were collected in 15-minute intervals up to 1-hour post-infection. Cells were then lysed in urea lysis buffer + 1% 2-mercaptoethanol and boiled for up to 5-minutes. Proteins were segregated by electrophoresis through a 4-12% Bis-Tris polyacrylamide gel (Invivogen), transferred to a PVDF membrane, and probed for IKKɑ/β, phospho-IKKɑ/β (serine 176/180), IκBɑ, and actin. Anti-rabbit or anti-mouse IgG conjugated to HRP were used as secondary antibodies. Blots were developed using SuperSignal West Femto Maximum Sensitivity Substrate (Thermo). All antibodies were purchased from Cell Signaling Technology.

### Microscopy

BMDMs were prepared as described and seeded into 8-well glass microscope slides (Millicell EZ slide, Millipore) and incubated overnight at 37°C. The following morning, the cells were infected as described with Δ*rgg2*, Δ*rgg3*, Δ*rgg2* + Δ*rgg3* GAS, or bacteria-free media at an MOI of 50. At sequential 15- minute intervals post-infection, cells were washed with PBS and fixed with 4% paraformaldehyde for 15 minutes. The cells were then washed with PBS three times then blocked with 5% BSA and 0.3% triton-X in PBS for 60m at room temperature or 4°C overnight. The blocking buffer then replaced with PBS containing 1% BSA, 0.3% triton-X and anti-p65 antibody (Cell Signaling Technology), then incubated at 4°C overnight. Cells were then washed and incubated with Alexa Fluor 488 anti-rabbit IgG (H+L) F(ab) 2 antibody (Cell Signaling Technology) for 2 hours at room temperature. 1g/mL of DAPI (Biotium) was added for 5 minutes then the cells were washed three times. Glass coverslips were mounted onto slides with Prolong Diamond Antifade Mountant (Invitrogen) and cured overnight at room temperature in the dark. Slides were visualized on an inverted fluorescent microscope (Keyence BZ-X710).

### NFκB Translocation Assays

Differentiated THP-1 were prepared in RPMI 1640 and infected as described above with Δ*rgg2*, Δ*rgg3*, Δ*rgg2* + Δ*rgg3* GAS, or bacteria-free media and collected after 35-minutes post-infection. Lysis, nuclear isolation, and the subsequent assay were performed according to kit manufacturer instruction (TransAM NFκB Family Kit from Active Motif). Nuclear protein input was normalized used a Bradford assay (Bio Rad)

### Pathway Inhibition and Induction

Pharmacological inhibitors were added either 2.5h prior to infection or the day before infection at the time of seeding. Inducers were added 30m prior to infection or the day before infection at the time of seeding. Cells were treated with the agent during the infection and through the time they were collected. Concentrations were as follows: 20 µM AKT1/2 inhibitor (Sigma A6730); 7.5 µM TWS119 (Sigma); 7.5 µM DIF-3 (MedChem Express); 10 µM SC79 (MedChem Express); 2 µM CDK2 inhibitor II (Cayman); 5 µM GSK126 (Selleck Chemicals); 500 nM Suberoylanilide hydroxamic acid (SAHA); 10 µM EX527 (Cayman); 5 µM AGK2 (Cayman); 1 µM BRM014 (Cayman).

### Griess Assay

70,000 RAW 264.7 cells were plated in a 96-well plate the day before infection. The following day, cells were infected with GAS at an MOI 10 as described above. Relative nitrite production by macrophages in response to Group A Strep was measured by Griess assay ∼18 hours post-infection according to manufacturer’s instructions (Promega).

### NFκB Reporter Assay

35,000 RAW Blue cells were plated in a 96-well plate the day before infection. The following day, cells were infected with GAS at an MOI 10 as described above. 150µL of Quantiblue Reagent (Invivogen) was mixed with 50µL of sample supernatant in a new 96-well plate, and the absorbance was read at 625nm every 30m for 6 hours.

### Reinfections

RAW 264.7 or Raw Blue cells were seeded and infected as described above in 6-well plates. The day after the infection, supernatant was collected, and the suppression phenotype was confirmed by Griess assay or Quantiblue assay. Cells were then collected and reseeded for a secondary infection 2 days post primary infection. Alternatively, the cells were allowed to rest for 2 days after the primary infection before reseeding. They were then infected for the secondary infection 4 days post primary infection.

### ELISA

RAW 264.7 or THP-1 cells were prepared and infected as described above. At appropriate timepoints described in the text, supernatants were collected, centrifuged, re-collected, and frozen at - 80°C. All ELISAs were performed with kits provided by BioLegend.

## Acknowledgements

We’d like to thank Jason Wood and Mark Maienschein-Cline at the UIC Research Informatic Core for their assistance with the clustering analysis of the RNAseq data, to Dr. Mat Wietecha for helpful discussions regarding the RNAseq and PTM analysis, and to Jennifer Chang and Ian McIntire for their generous technical assistance and general discussions.

This study was supported by NIH R01AI162679-01A1 to M.J.F and NIH F31AI147429 to K.M.R.

## Data Availability

The raw and processed data from the PTM scan can be found in the MassIVE repository. doi:10.25345/C51834F1P.

Raw RNAseq data will be available on Gene Expression Omnibus at a later date.

## Supplemental

**Figure S1.**
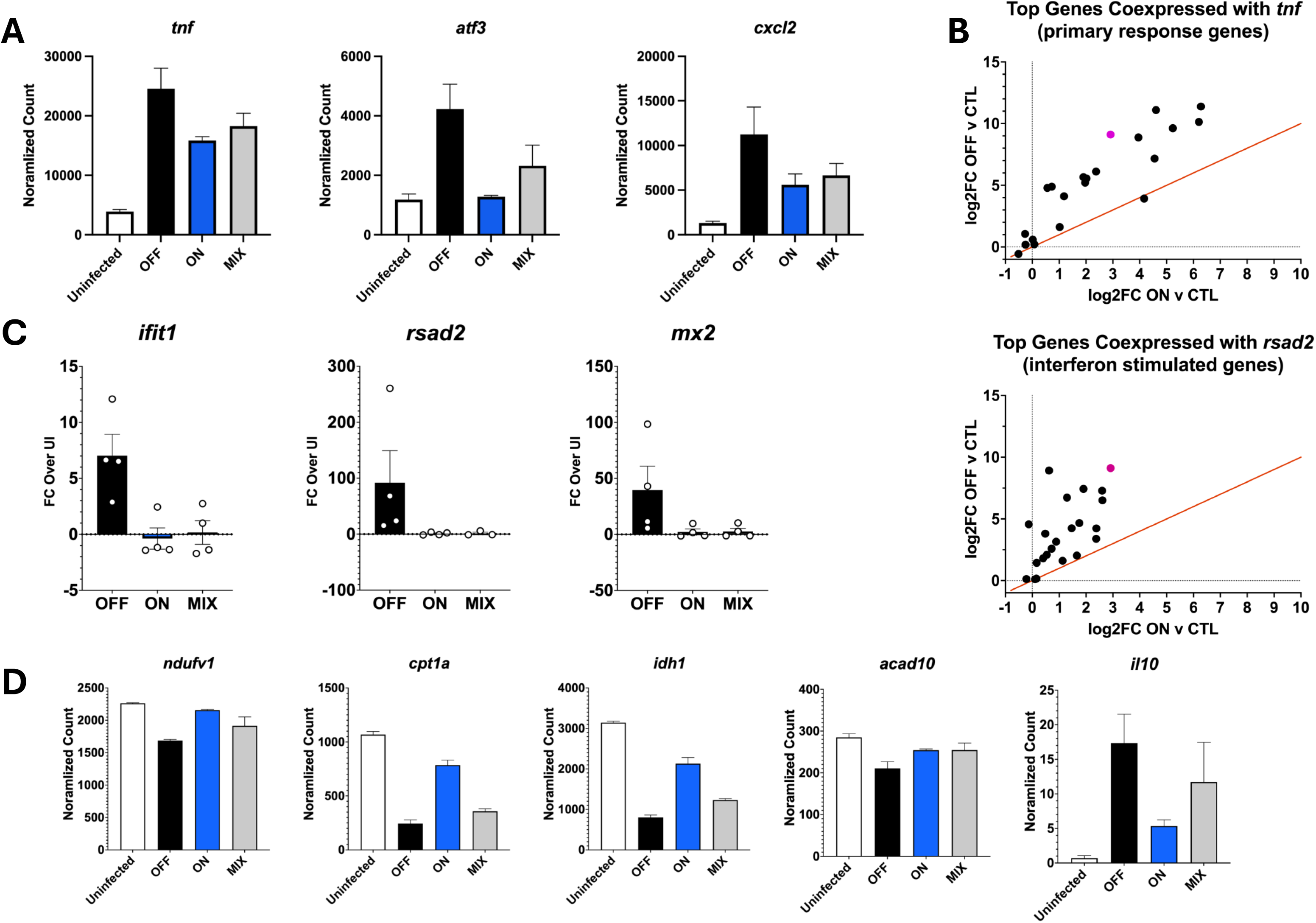
A) Normalized read counts of infected RAW 264.7 cells at 2h post-infection (p.i.) of select early response genes from RNA-seq. B) Comparison of log_2_ fold changes of QS-OFF-infected over uninfected samples, and to QS-ON-infected over uninfected samples, averaged across all time points from RNA-seq dataset for genes most commonly co-expressed with *tnf* and *rsad2*. Genes most commonly co-expressed with these genes were determined using ARCHS4’s RNA-seq gene-gene co-expression matrix acessed via Enrichr. Orange line shows the line of identity. C) RT-qPCR of select type I interferon stimulated genes showing fold change over uninfected cells at 4h p.i. D) Normalized read counts at 8h p.i. from RNA-seq dataset of select genes associated with oxidative phosphorylation and fatty acid metabolism, and *il10*.

**Figure S2.**
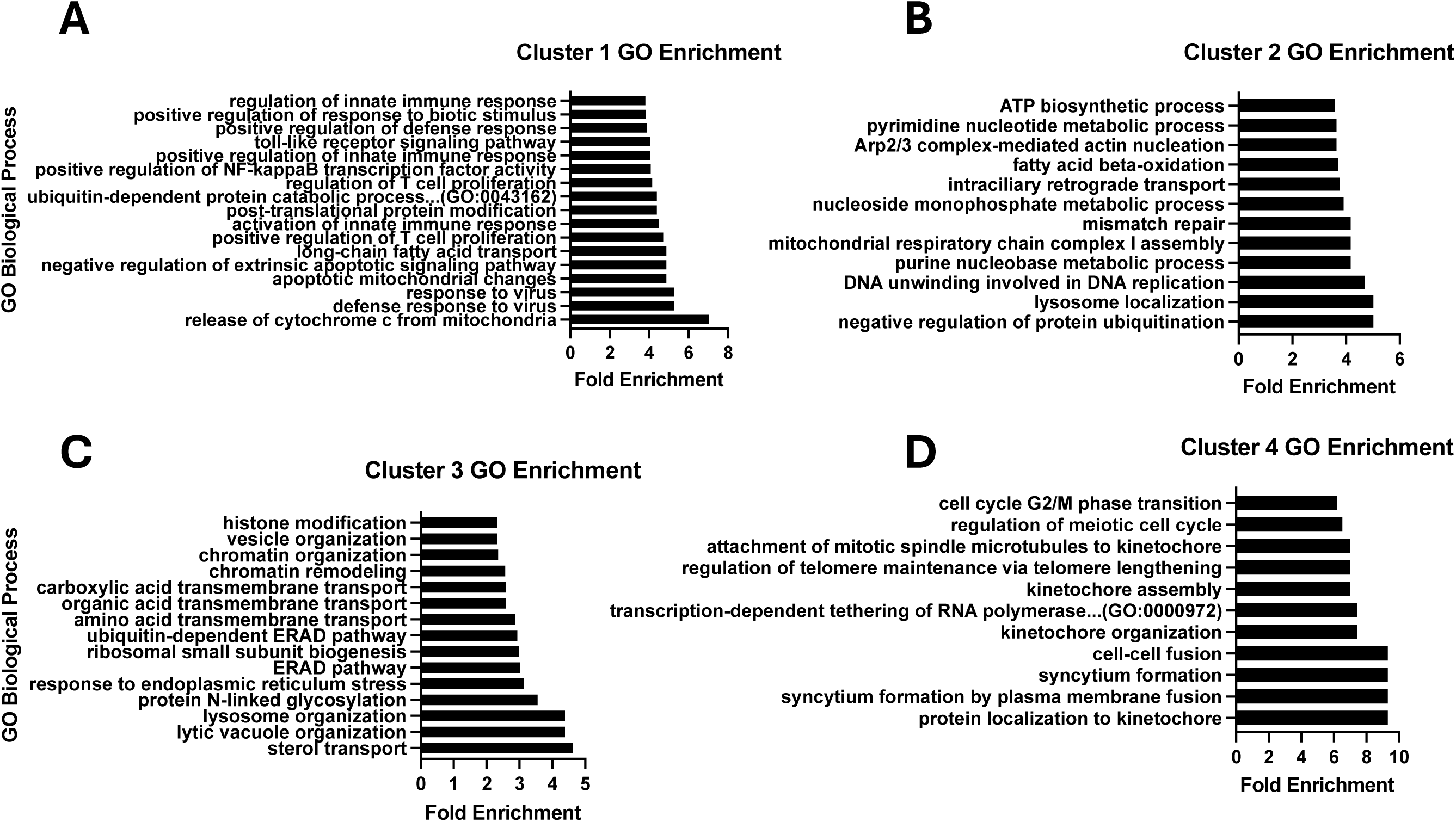
Enrichment of gene ontology (GO) biological processes clusters I-IV (A-D, respectively) created from analysis of RNA-seq dataset at 8h post-infection.

**Figure S3.**
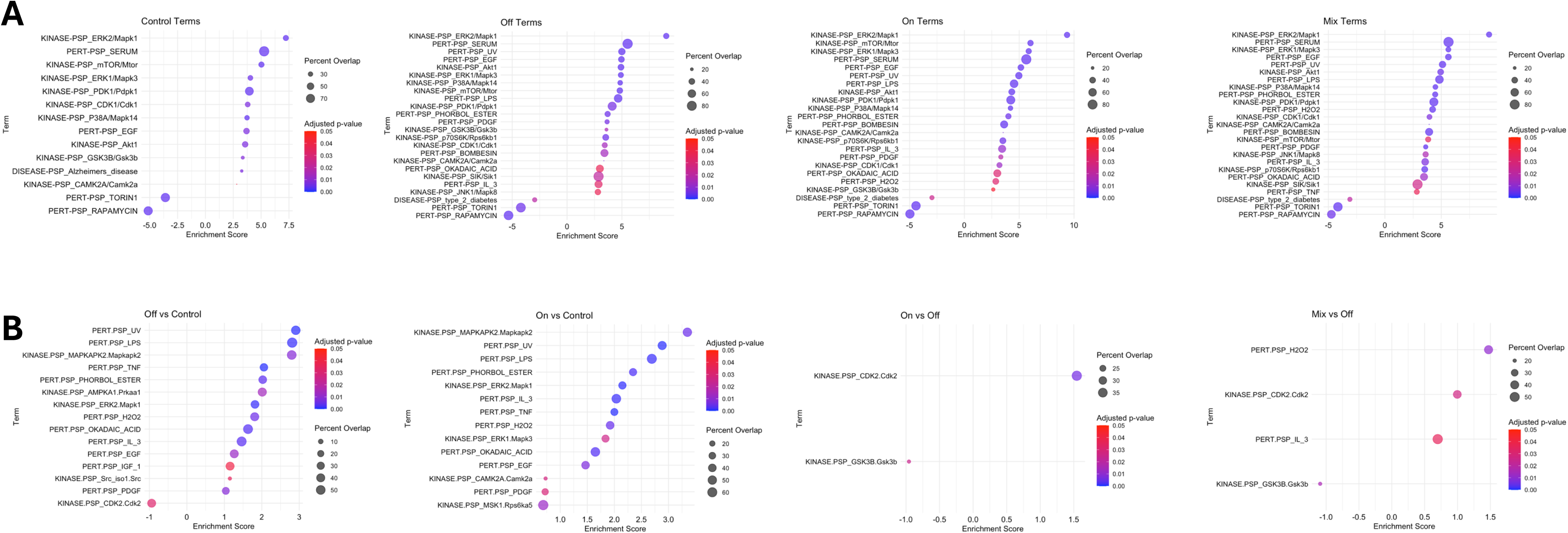
A) PTM-SEA pathways that are significantly enriched (fdr-adjusted p-value<.05) from the post translational modification scan of each infection condition B) PTM-SEA pathways that show significant differences in enrichments scores (p-value<0.05) in select pairwise comparisons. Significance determined by one way ANOVA.

**Supplemental Table 1.**
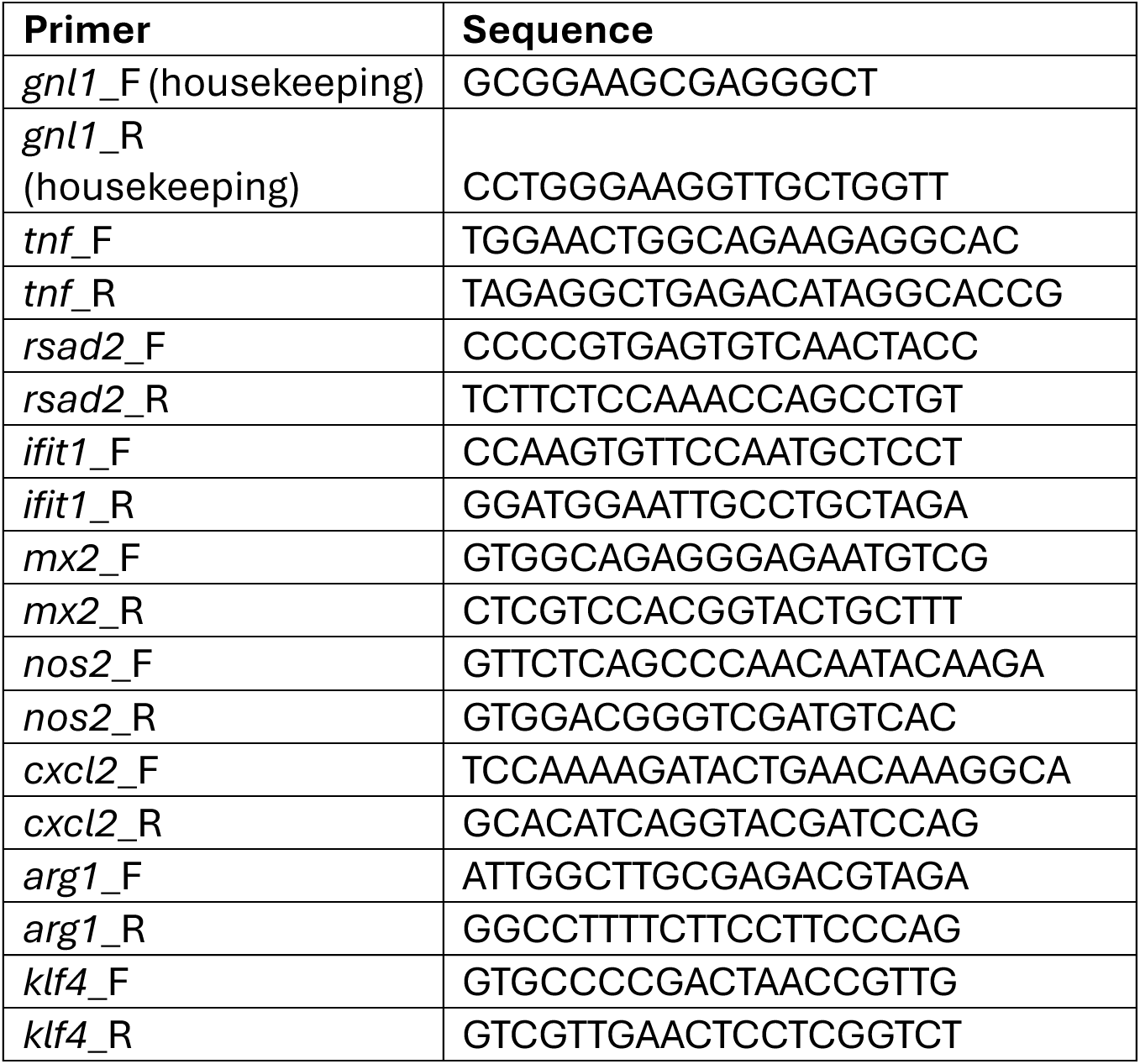
Primers used for RT-qPCR.

